# Associations Between Resting State Functional Connectivity and a Hierarchical Dimensional Structure of Psychopathology in Middle Childhood

**DOI:** 10.1101/2020.04.28.065086

**Authors:** Nicole R. Karcher, Giorgia Michelini, Roman Kotov, Deanna M. Barch

## Abstract

**Background:** Previous research from the Adolescent Brain Cognitive Development (ABCD) study delineated and validated a hierarchical 5-factor structure with a general psychopathology (‘p’) factor at the apex and five specific factors (internalizing, somatoform, detachment, neurodevelopmental, externalizing) using parent-reported child symptoms. The current study is the first examining associations between dimensions from a hierarchical structure and resting state functional connectivity (RSFC) networks.

**Methods:** Using 9-11-year-old children from the ABCD baseline sample, we compared the variance explained by each hierarchy level (p-factor, 2-factor, 3-factor, 4-factor, and 5-factor models) in RSFC. Analyses were first conducted in a discovery dataset (n=3790) with significant associations examined in a replication dataset (n=3791).

**Results:** The current study found associations between p-factor and lower connectivity within default mode network (DMN), although stronger effects emerged for the neurodevelopmental factor. Neurodevelopmental impairments were related to variation in RSFC networks associated with attention to internal states and external stimuli. These networks included within DMN, DMN with cingulo-opercular (CON) and ‘Other’ (Unassigned) networks, CON with ventral attention and ‘Other’ network, and dorsal attention with ‘Other’ network. Results held when accounting for parental psychopathology.

**Conclusion:** The hierarchical structure of psychopathology showed replicable links to RSFC alterations in middle childhood. The p-factor had minimal association with altered connectivity, while the specific neurodevelopmental dimension showed robust associations with multiple RSFC impairments. Results show the utility of examining associations between intrinsic brain architecture and specific dimensions of psychopathology, revealing associations specifically with neurodevelopmental impairments.

---

The Research Domains Criteria (RDoC)^1,2^ and Hierarchical Taxonomy of Psychopathology (HiTOP)^3,4^ are transdiagnostic initiatives that have pushed psychiatric research to move beyond traditional diagnostic categories and to examine psychiatric problems as a multidimensional structure. One advantage of this approach is that dimensional constructs may align more closely to underlying neural mechanisms than diagnoses.^2,5^ The HiTOP model proposes that dimensions of psychopathology are organized hierarchically, from narrowest to broadest, with each dimension potentially providing important information in terms of functional and biological correlates.^6,7^

Several dimensions of psychopathology have been identified in children and adolescents.^8–11^ In particular, a study of the Child Behavioral Checklist (CBCL)^8^ in Adolescent Brain Cognitive Development (ABCD) data found a hierarchical structure of psychopathology with a broad general “p-factor” at the apex, which progressively differentiates into narrower factors, with the most fine grained structure representing five lower-order major dimensions (internalizing, externalizing, neurodevelopmental, somatoform, and detachment).^12^ The p-factor, also known as a general psychopathology factor,^13–16^ has been hypothesized as critical to understanding mental disorders.^13^ Alternatively, others suggested that p-factor may be too general and heterogeneous to reveal etiology, potentially representing functional consequences of psychopathology.^17,18^ Hence, it remains unclear what level of specificity in phenotypes is most informative for understanding neural mechanisms.

At other levels of the hierarchy,^12^ the 2-factor solution was comprised of broad internalizing and broad externalizing factors consistent with prior research.^19–21^ In the 3-factor structure, a neurodevelopmental factor (e.g. inattention, hyperactivity, impulsivity, clumsiness, repetitive behaviors) emerged from broad externalizing and internalizing dimensions. For the 4-factor solution, somatoform problems separated from the broad internalizing factor. In the 5-factor structure, the broad internalizing factor split into internalizing problems (e.g. anxiety, some mood symptoms) and detachment (e.g. social withdrawal). In this model, internalizing, externalizing, detachment, and somatoform map clearly on the corresponding HiTOP spectra. Although a neurodevelopmental spectrum is not yet represented in HiTOP, this ABCD study and other evidence supports its validity^22^ and potential inclusion of a neurodevelopmental spectrum in dimensional models of psychopathology. A similar hierarchical structure emerged for self-reported symptoms in the parents.

This hierarchical structure was also validated using a number of clinically relevant measures. While the child p-factor alone was sufficient to account for some clinical variables (e.g., medical and mental health service utilization), more fine grained dimensions, including some from the 5-factor structure, were necessary to more fully account for variance in other clinically relevant features, such as developmental delays, cognitive, social and educational functioning. Overall, this previous study delineated a hierarchical dimensional structure of psychopathology in one of the largest samples of children available to date, but also provided evidence for the incremental clinical utility of levels in this hierarchy. The current study sought to expand upon this previous investigation to examine the associations between the previously identified hierarchical structure of psychopathology and resting state functional connectivity (RSFC) in the ABCD study.

Previous studies in children and adolescents have examined the neural correlates of psychopathology, including RSFC. RSFC is based on using the temporal correlations of spontaneous fluctuations in BOLD to parse the brain into functionally organized networks of brain regions.^23^ RSFC can be particularly useful for understanding brain-behavior relationships, as it can be used to examine the entire functional architecture of the brain, has low participant burden, and there is evidence that the findings can be reproducible.^24^ Further, alterations in brain network organization during development are implicated in the emergence of psychopathology.^25^

Previous RSFC studies either focused on a limited set of psychopathology dimensions (e.g. internalizing and externalizing) or on diagnostic categories,^26–28^ and did not examine hierarchy of psychopathology dimensions consistent with the HiTOP model. Among children and adolescents, greater p-factor scores derived from the CBCL were associated with reduced maturation of the default mode network (DMN), although this was driven primary by neurodevelopmental symptoms.^29^ The DMN is a group of functionally correlated brain regions showing lower activation during goal-oriented tasks^30^ and is involved in attention to internal states.^23^ Externalizing problems, including aggression and risky behaviors, and neurodevelopmental symptoms, including attention-deficit/hyperactivity disorder (ADHD) symptoms, have also been associated with a number of connectivity alterations.^22,27^ These include lower anticorrelation (negative RSFC) between DMN and both cingulo-opercular (CON; a network associated with information integration, including salience attribution^31^)^32^ and sensorimotor regions.^27^ The externalizing dimension was also linked to alterations in connectivity with the salience network (network involved in detection of relevant stimuli), such as lower connectivity with the DMN.^27,33^ Other studies found that child and adolescent internalizing symptoms (i.e., encompassing symptoms of anxiety and depression) were associated with alterations in connectivity in DMN regions,^27,33,34^ as well as disruptions in the Ventral Attention Network (VAN; a network associated with orienting and responding to novel stimuli^35^).^33,36^

Despite these promising findings, several questions remain unanswered regarding the associations between psychopathology and RSFC in middle childhood. First, most previous studies only delineated a general factor and/or internalizing and externalizing dimensions, despite evidence that more specific dimensions, such as a neurodevelopmental spectrum, may be important for investigating associations with clinically relevant risk indicators and outcomes.^12^ No previous research has investigated the associations between RSFC and hierarchically-organized psychopathology dimensions consistent with the HiTOP model. Second, it is unclear whether alterations in connectivity are associated with specific psychopathology dimensions over and above a general psychopathology p-factor. Third, although previous studies were typically based on parent-reported child psychopathology, no previous research has examined whether associations between parent-rated child psychopathology and RSFC may be explained by parental psychopathology, which may plausibly occur due to factors such as genetic or shared environmental influences.

The current study will address these gaps by examining the relationship between middle childhood parent-reported hierarchical dimensions of psychopathology and RSFC throughout the brain (13 networks from the “Gordon” parcellation) in a large sample of 9-11-years-old from the ABCD study. Based on previous research, we hypothesized that the p-factor, internalizing, and neurodevelopmental symptoms would be associated with alterations in DMN connectivity, and that internalizing symptoms would be associated with VAN connectivity alterations. We tested these specific hypotheses in the context of an unbiased “whole brain network parcellation” approach. The current study also importantly incorporated best analytic practices, including examining results in a discovery dataset and then testing findings in a replication dataset. We used hierarchical multiple regression to test whether finer-grained factors account for variance over and above p-factor when examining within- and between-network RSFC.^37–39^ Further, all analyses were followed-up by examining whether results held when accounting for parental psychopathology.

## Methods

### Participants

A sample of 11,873 individuals was obtained from the ABCD study, a large-scale study tracking 9-11-years-olds recruited from 21 research sites across the United States.^40^ Potential participants were excluded from ABCD study participation for the following reasons: child not fluent in English, MRI contraindication (e.g., irremovable ferromagnetic implants or dental appliances, claustrophobia, pregnant), major neurological disorder, gestational age less than 28 weeks or birthweight less than 1,200 grams, history of traumatic brain injury, or current diagnosis of schizophrenia, autism spectrum disorder (moderate, severe), mental retardation/intellectual disability, or alcohol/substance use disorder. These data were accessed from the National Institutes of Mental Health Data Archive (DOI 10.15154/1460410; see Acknowledgments). The current study is based on 9,987 unrelated children (randomly selecting one child per family when more than one participated) from the baseline ABCD 2.0.1 data release.^12^ Participants were removed from analyses in the current study either for not having at least one resting state scan that passed quality assurance criteria (n=550) or being run on a Philips scanner (n=1208) due to an error in processing in the 2.01 ABCD data release, or in the case of missing data (n=631; Supplemental Table 1). Next, the remaining dataset was divided in discovery (n=3790) and replication (n=3791) datasets (Table 1).

**Table 1.** Demographic Characteristics for the Discovery and Replication Datasets^a^

|  | Discovery Dataset (n=3790) | Replication Dataset (n=3791) | Test Statistic<sup>b</sup> | p value |
| --- | --- | --- | --- | --- |
| Sex (% Female) | 48.9% | 47.5% | 1.349 | .25 |
| Race/Ethnicity (% White) | 50.8% | 51.3% | .941 | .92 |
| Age (in months) | 119.040 (7.445) | 118.780 (7.412) | -1.191 | .23 |
| Financial Adversity | 0.476 (1.111) | 0.485 (1.122) | 0.386 | .67 |
| Average Head Motion | 0.250 (0.229) | 0.257 (0.238) | 1.322 | .19 |
| Scanner type (% Siemens) | 70.3% | 72.0% | 2.739 | .10 |
| Hierarchical Dimensions of Psychopathology |  |  |  |  |
| P-factor | 0.020 (0.944) | 0.026 (0.915) | 0.323 | .75 |
| 2-factor |  |  |  |  |
| Internalizing | 0.041 (0.900) | 0.035 (0.896) | -0.310 | .76 |
| Externalizing | 0.033 (0.891) | 0.043 (0.864) | 0.495 | .62 |
| 3-factor |  |  |  |  |
| Internalizing | 0.060 (0.893) | 0.049 (0.886) | -0.495 | .62 |
| Externalizing | 0.049 (0.850) | 0.054 (0.829) | 0.243 | .81 |
| Neurodevelopmental | 0.020 (0.830) | 0.034 (0.833) | 0.733 | .46 |
| 4-factor |  |  |  |  |
| Internalizing | 0.055 (0.855) | 0.058 (0.831) | 0.15 | .88 |
| Externalizing | 0.061 (0.860) | 0.049 (0.861) | -0.604 | .55 |
| Neurodevelopmental | 0.028 (0.831) | 0.044 (0.830) | 0.842 | .40 |
| Somatoform | 0.069 (0.779) | 0.065 (0.774) | -0.24 | .81 |
| 5-factor |  |  |  |  |
| Internalizing | 0.065 (0.867) | 0.067 (0.838) | 0.079 | .94 |
| Externalizing | 0.041 (0.841) | 0.051 (0.828) | -0.713 | .48 |
| Neurodevelopmental | 0.041 (0.841) | 0.057 (0.832) | 0.815 | .42 |
| Somatoform | 0.093 (0.790) | 0.085 (0.781) | -0.443 | .66 |
| Detachment | 0.049 (0.749) | 0.039 (0.761) | -0.595 | .55 |

<sup>a</sup>Means (SDs) or percentages (as indicated).<sup>b</sup>Independent-samples *t* tests were used to compare means for the exposed and unexposed groups. $\chi^2$ tests were used to compare ordinal/binary variables across groups.

### Measures

We used psychopathology dimensions factor-analytically derived in ABCD^12^ using the parent-rated CBCL^21^ and the self-reported Adult Self Report (ASR)^41^ from ASEBA. Parents (mean age=39.94, SD=6.93; 89.03% female) rated their own (ASR) and their child’s (CBLC) psychopathology occurring in the past 6 months on 3-point scale (i.e., 0=never, 1=sometimes, 2=often). ASR factors were included to examine whether RSFC results held when accounting for parental psychopathology. The hierarchical structure used in this study was delineated in the previous investigation through exploratory factor analysis with oblique (goemin) rotation, whereby the maximal number of factors was determined using parallel analyses and interpretability of factor solutions.^42^ For more details of the creation of factor scores used in analyses, see (Michelini et al.,2019).^12^

### Imaging Procedure

ABCD imaging procedures have been detailed in previous studies.^43,44^ All children were run on a 3T scanner (either Siemens or General Electric) with a 32-channel head coil and completed T1-weighted and T2-weighted structural scans (1mm isotropic). Participants also completed four 5-minute resting-state BOLD scans, with their eyes open and fixated on a crosshair. Resting state images were acquired in the axial plane using an EPI sequence. Other resting-state image parameters varied by 3T scanner and have been previously detailed (https://abcdstudy.org/images/Protocol_Imaging_Sequences.pdf). A Fisher Z-transformed average of all pair-wise correlations within each of the 13 Gordon networks (e.g., within the DMN or FPN) and between each of the 13 networks with the other 12 networks (e.g., between the DMN and the FPN) were examined.^37^

### Statistical analysis

We first randomly split the data into discovery (n=3790) and replication (n=3791) datasets. All analyses were first conducted in the discovery dataset. Every model was conducted as a hierarchical linear model (HLMs) using the R lme4 package.^45^ The research sites were modeled as random intercepts (to account for nonindependence of observations) and every model included age, sex, financial adversity, race/ethnicity, average motion, and scanner type as covariates. Analyses proceeded in three steps.

#### Step 1

Following the general approach of the previous study delineating and validating the investigated hierarchical structure^12^, we entered factor structures with a progressively greater number of factors (baseline model [only covariates]; p-factor; 2-factor: internalizing and externalizing; 3-factor: internalizing, externalizing, neurodevelopmental; 4-factor: internalizing, externalizing, neurodevelopmental, somatoform; 5-factor: internalizing, externalizing, neurodevelopmental, somatoform, detachment) as blocks of HLMs with each RSFC metric as dependent variables. Specifically, using HLMs we tested for every outcome metric whether each hierarchy level as a block provided a significantly better fit than all levels above it (e.g., whether the 5-factor model outperformed 4-, 3-, 2-, and 1-factor models as well as the covariates). Changes in fit were assessed using change in R^2^ [ΔR^2^] and chi-square tests. Any factor structures that passed False Discovery Rate (FDR) correction in the discovery dataset were examined in the replication dataset.

#### Step 2

When adding a factor structure produced a FDR-corrected significant increase in variance explained (i.e., FDR *p*<.05 with 65 FDR-corrected comparisons for each network [5 models x 13 networks]), we ran an additional HLM to examine the association with each of the individual dimensions included in that factor structure in both the discovery and replication datasets. For example, if the 5-factor structure accounted for a significant increase in variance over simpler structures, we examined an additional model only including the significant structure (e.g., 5-factor structure) to examine *which* of the dimensions in the 5-factor model was responsible for the increment in variance accounted for.

#### Step 3

For any dimension that was statistically significant in Step 2, a HLM examined the association between that structure (e.g., 5-factor structure) and the RSFC metric including parental psychopathology in the discovery and replication datasets. In these models, for any dimension that was statistically significant (e.g., child neurodevelopmental scores) we included the corresponding parental psychopathology dimension (e.g., parental neurodevelopmental scores).

We used the HLM procedure described above to analyze the associations between factor scores (i.e., 1(p)-factor, 2-factor, 3-factor, 4-factor, and 5-factor extracted factor scores as in previous research^12^) and functional connectivity within and between each of the 13 Gordon networks^37^ [Auditory (AUD), Cingulo-Opercular (CON), Cingulo-Parietal (CPAR), Default Mode (DMN), Dorsal Attention (DAN), Fronto-Parietal (FPN), Other (also referred to as the “Unassigned” network), Retrosplenial-Temporal (RSP), Salience (SA), Sensorimotor-Hand (SMH), Sensorimotor-Mouth (SMM), Ventral Attention (VAN), Visual (VS)]. See Supplemental Table 2 for overall means for each of the within- and between-network metrics. Results are expressed as standardized estimates (*β*s).

## Results

### Associations between RSFC and the Hierarchical Dimensions of Psychopathology

#### 1(p)-factor model

As predicted, lower within-network connectivity in the DMN was significantly associated with higher child p-factor scores. Specifically, the 1(p)-factor model (Table 2 and Figure 1) accounted for a significant increment in variance over the baseline model. Thus, the p-factor was significantly associated with lower within-network DMN connectivity, *χ*^2^(1)=11.19, *p*<.001 (see Table 3 and Figure 2; see Supplemental Figure 1 for scatterplots). There were no other RSFC associations with child p-factors scores that passed FDR in the discovery dataset and replicated in the replication dataset.

**Figure 1.**
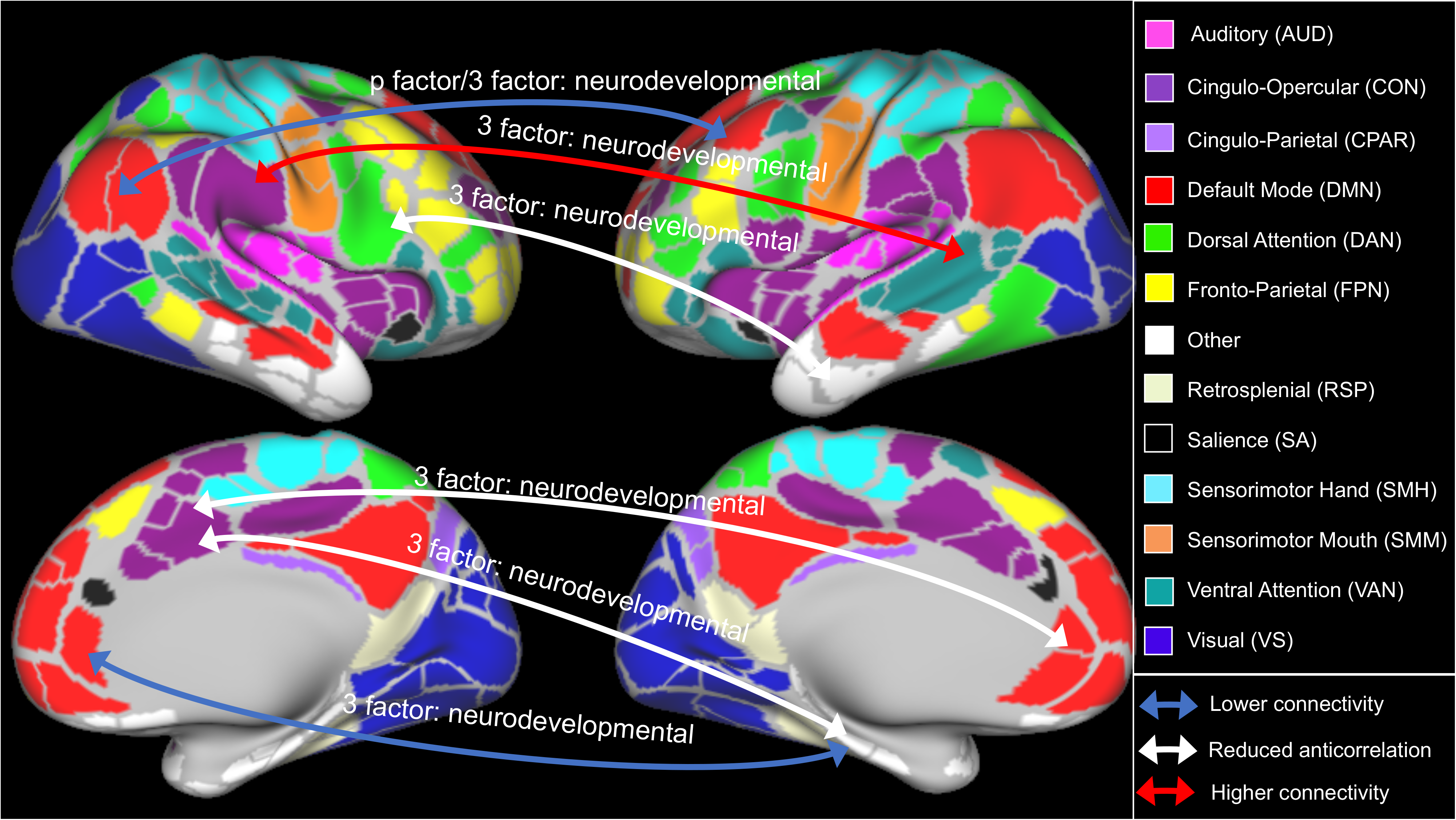
Proportion of variance explained (R^2^) for each factor model (1- to 5-factor solutions) for with resting state functional connectivity metrics in the discovery dataset. Asterisks indicate significant change in R^2^ for that factor model versus the more parsimonious model (\**p*<.05, \*\**p*<.01, \*\*\**p*<.001).

**Figure 2.**
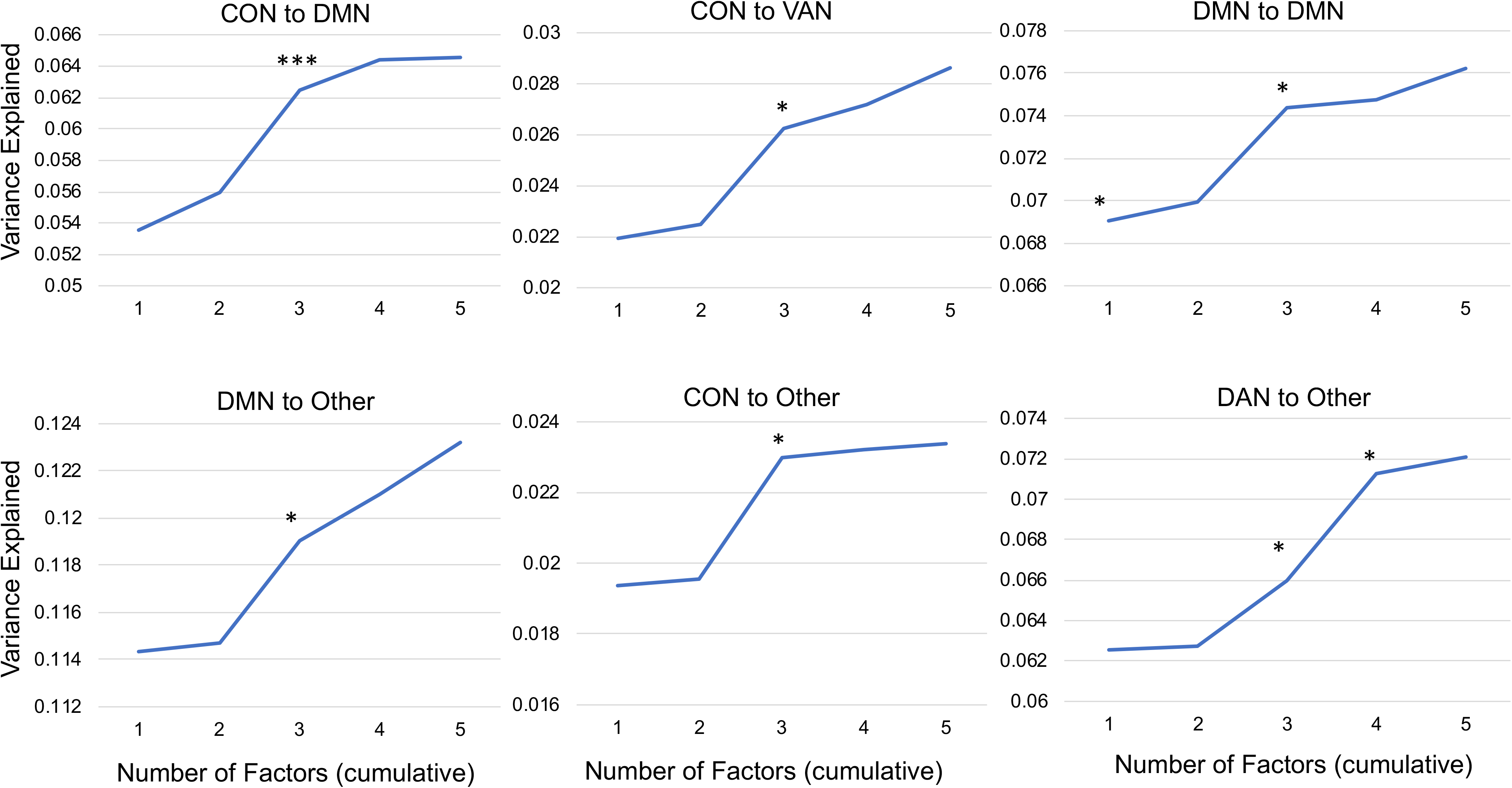
An illustration of all significant RSFC associations with hierarchical dimensions of psychopathology for the Gordon network parcellation.

**Table 2.** Proportion of Variance Explained in the Discovery and Replication Sets for All Models Surviving False Discovery Rate (FDR) correction in the Discovery Set^a^

| RSFC Metric | Model | Discovery Set |  |  | Replication Set |  |  |
| --- | --- | --- | --- | --- | --- | --- | --- |
| | | $\chi^2$ | FDR $p$ | $\Delta R^2$ | $\chi^2$ | $p$ | $\Delta R^2$ |
| CON to CON | 3 factor | 23.986 | <.001 | 0.59% | 4.369 | .22 | 0.13% |
| **CON to DMN** | **3 factor** | **27.489** | **<.001** | **0.66%** | **16.927** | **.001** | **0.44%** |
| **CON to VAN** | **3 factor** | **14.893** | **.04** | **0.38%** | **9.971** | **.02** | **0.24%** |
| DMN to AUD | 3 factor | 19.492 | .01 | 0.47% | 2.234 | .53 | 0.05% |
| **DMN to DMN** | **p-factor** | **11.192** | **.01** | **0.25%** | **7.889** | **.01** | **0.17%** |
| **DMN to DMN** | **3 factor** | **18.833** | **.01** | **0.45%** | **19.648** | **<.001** | **0.51%** |
| **DMN to Other** | **3 factor** | **18.415** | **.01** | **0.43%** | **9.352** | **.03** | **0.23%** |
| DMN to RSP | 3 factor | 13.005 | .04 | 0.33% | 1.505 | .68 | 0.04% |
| DMN to SA | 5 factor | 17.287 | .04 | 0.44% | 4.076 | .54 | 0.10% |
| DMN to VAN | p-factor | 10.423 | .006 | 0.30% | 2.372 | .12 | 0.06% |
| FPN to FPN | 4 factor | 18.279 | .04 | 0.45% | 4.876 | .30 | 0.12% |
| FPN to Other | 4 factor | 18.438 | .04 | 0.47% | 7.747 | .10 | 0.20% |
| **CON to Other** | **3 factor** | **13.510** | **.04** | **0.34%** | **18.707** | **<.001** | **0.49%** |
| **Other to DAN** | **3 factor** | **15.981** | **.01** | **0.32%** | **7.415** | **.03** | **0.19%** |
| Other to DAN | 4 factor | 18.981 | .01 | 0.53% | 2.055 | .84 | 0.04% |
| Other to RSP | 5 factor | 22.338 | .01 | 0.58% | 8.541 | .13 | 0.22% |
| SA to SA | 5 factor | 21.596 | .04 | 0.01% | 7.447 | .19 | 0.19% |

Abbreviations: RSFC=resting state function connectivity; p=p-value; $R^2$ =marginal proportion of variance explained by the entire model; $\Delta R^2$ =change in $R^2$ from the more parsimonious model; AUD=Auditory Network; CON=Cingulo-opercular Network; DMN=Default Mode Network; DAN=Dorsal Attention Network; FPN= Fronto-parietal Network; RSP= Retrosplenial Network; SA=Saliency Network; VAN=Ventral Attention Network.
<sup>a</sup>Models that are significant in both the Discovery and Replication sets are in bold. FDR-correction was only applied to the Discovery dataset, not the Replication dataset.

**Table 3.** Model Estimates for all Dimensions for Significant Models Examining Associations between Resting State Functional Connectivity (RSFC) and Hierarchical Dimensions of Psychopathology^a^

|  | Discovery Set |  |  | Replication Set |  |  |
| --- | --- | --- | --- | --- | --- | --- |
| | $\beta$ | $t$ | $p$ | $\beta$ | $t$ | $p$ |
| P-factor |  |  |  |  |  |  |
| DMN to DMN | -0.055 | -3.350 | .001 | -0.045 | -2.823 | .005 |
| 3-Factor |  |  |  |  |  |  |
| CON to DMN |  |  |  |  |  |  |
| Internalizing | -0.104 | -4.531 | <.001<sup>b</sup> | -0.050 | -2.333 | .02 |
| Externalizing | 0.012 | 0.593 | .55 | 0.012 | 0.638 | .52 |
| Neurodevelopmental | 0.125 | 5.637 | <.001 | 0.099 | 4.752 | <.001 |
| CON to VAN |  |  |  |  |  |  |
| Internalizing | -0.025 | -1.055 | .29 | -0.031 | -1.378 | .17 |
| Externalizing | 0.032 | 1.520 | .13 | 0.006 | 0.321 | .75 |
| Neurodevelopmental | 0.046 | 2.012 | .04 | 0.068 | 3.109 | .002 |
| DMN to DMN |  |  |  |  |  |  |
| Internalizing | 0.052 | 2.290 | .02 | 0.019 | 0.857 | .39 |
| Externalizing | -0.012 | -0.589 | .56 | 0.023 | 1.223 | .22 |
| Neurodevelopmental | -0.114 | -5.150 | <.001 | -0.104 | -4.894 | <.001 |
| DMN to Other |  |  |  |  |  |  |
| Internalizing | 0.049 | 2.174 | .03 | 0.005 | 0.242 | .81 |
| Externalizing | 0.003 | 0.173 | .86 | 0.028 | 1.536 | .13 |
| Neurodevelopmental | -0.088 | -4.101 | <.001 | -0.064 | -3.095 | .002 |
| CON to Other |  |  |  |  |  |  |
| Internalizing | -0.060 | -2.271 | .02<sup>b</sup> | -0.071 | -2.818 | .005 |
| Externalizing | -0.022 | -0.920 | .36 | 0.005 | 0.223 | .82 |
| Neurodevelopmental | 0.099 | 3.652 | <.001 | 0.119 | 4.563 | <.001 |
| DAN to Other |  |  |  |  |  |  |
| Internalizing | -0.059 | -2.553 | .01 | -0.008 | -0.362 | .72 |
| Externalizing | -0.022 | -1.067 | .29 | -0.023 | -1.202 | .23 |
| Neurodevelopmental | 0.083 | 3.743 | <.001 | 0.057 | 2.654 | .008 |

Abbreviations: $\beta$ =standardized regression coefficient; $t$ =t-test test statistic; FDR=False Discovery Rate; $p$ =p-value; DMN=Default Mode Network; CON=Cingulo-opercular Network; DAN=Dorsal Attention Network; VAN=Ventral Attention Network.
<sup>a</sup>Models with connectivity as the outcomes and dimensions entering simultaneously as IVs, and controlling for age, sex, race/ethnicity, financial adversity, average head motion, and scanner type as covariates
<sup>b</sup>Although these models are significant in discovery and replication samples, when examining a model with just internalizing symptoms with RSFC, internalizing symptoms is not significant, indicating multicollinearity.

#### 2-factor model

The 2-factor model (internalizing and externalizing) did not account for a significant increment in variance of any of the RSFC metrics over the model with covariates + p-factor in the discovery dataset at FDR threshold (Supplemental Tables 3-15).

#### 3-factor model

As can be seen in Table 2 and Figure 1, the 3-factor model (internalizing, externalizing, and neurodevelopmental) significantly accounted for additional variance over the baseline + p-factor + 2-factor model for several RSFC metrics, including within DMN, DMN to CON and “Other”, CON to VAN and “Other,” and DAN to “Other”. For within-network DMN connectivity, the follow-up analyses examining each individual factor of the 3-factor model indicated that lower within-network DMN connectivity was significantly associated with higher 3-factor neurodevelopmental factor scores (Table 3). In addition, follow-up analyses indicated that lower connectivity between the DMN and ‘Other’ networks was also associated with higher 3-factor neurodevelopmental factor scores (Table 3). Follow-up analyses further indicated that lower anticorrelation (i.e., less negative connectivity) between the CON and DMN, between the CON and ‘Other’, and between the DAN and ‘Other’ networks were associated with higher neurodevelopmental factor scores (Table 3). Lastly, follow-up analyses indicated greater connectivity between the CON and VAN was associated with greater neurodevelopmental scores. See Figure 2 for a summary of these results.

#### 4- and 5-factor models

The 4-factor (internalizing, externalizing, neurodevelopmental, and somatoform) and 5-factor (internalizing, externalizing, neurodevelopmental, somatoform, and detachment) models did not add a significant increment in RSFC variance over the simpler structures in both the discovery (and survived FDR correction) and replication datasets (Table 2, Supplemental Tables 3-15). Although for DAN and ‘Other’ connectivity, the 4-factor model accounted for additional variance over the baseline + p-factor + 2-factor + 3-factor in the discovery dataset, the association did not reach statistical significance in the replication dataset (*p*=.84, Table 2).

#### Controlling for Parental Psychopathology

Interestingly, the child p-factor was only nominally associated with lower DMN connectivity (child p-factor: *p*=.05; Supplemental Table 16) when including parental p-factor scores, though parental p-factor scores themselves were not related to lower DMN connectivity (parent p-factor: *p*=.22; Supplemental Table 16). Unlike the p-factor, the results for almost all of these 3-factor models remained significant when including the corresponding parental neurodevelopmental factor scores (*p*s<.005; Supplemental Table 16). The only exception was the association between neurodevelopmental factor and DAN to ‘Other’ connectivity, which became trend-level when parental neurodevelopmental scores where included (*p*=.06). In contrast, parental neurodevelopmental factor scores were not significantly associated with these RSFC metrics (*p*s>.15).

## Discussion

The current study contributes significantly to our understanding of the neural correlates of hierarchically-organized psychopathology dimensions consistent with the HiTOP model in middle childhood. The results provide novel information about the relationship of functional networks across the entire brain architecture to specific dimensions of psychopathology. Using a robust and rigorous 2-sample strategy, the current study tested hypotheses about the associations between RSFC and psychopathology dimensions in the ABCD baseline sample using an unbiased “whole brain networks” approach. Overall, we found evidence that greater neurodevelopmental problems were associated with within-network DMN under-connectivity, but also alterations in RSFC among other networks, including the CON, DAN, and ‘Other’ networks. There was some evidence for associations between p-factor and within-network DMN under-connectivity, albeit weaker than its link to the neurodevelopmental spectrum. Importantly, many of these associations remained when accounting for parental psychopathology. The current study indicates that specifically neurodevelopmental problems are associated with alterations in RSFC networks associated with attention to internal states and external stimuli (e.g., DMN, CON), which may have important implications for the etiology of these problems.

The current study found that higher p-factor was associated with only one RSFC marker, lower within-network DMN connectivity. This is consistent with previous findings of an association between DMN connectivity and general psychopathology.^29^ However, this association was no longer significant when accounting for parental p-factor scores. One possible explanation is that this pattern reflects evidence that the p-factor is heritable^46^ and thus child p-factor pathology may be closely related to parental p-factor pathology, and therefore it is this heritable pathology (as opposed to the child’s unique pathology) that is associated with RSFC network alterations. However, an alternative possibility is that this finding reflects the parents own mental health concerns coloring their rating of their child’s mental health. This would potentially indicate that greater parental psychopathology has downstream effects for the child’s RSFC. Moreover, the association between p-factor and lower DMN connectivity may be due to neurodevelopmental problems included in the p-factor. Overall, our results provide only minimal support for hypothesis that focus on general psychopathology would reveal robust neural signal. Instead, present findings suggest that p-factor is too heterogeneous to capture neural impairments, at least as reflected in RSFC networks.

In addition, the current study indicates that several early emerging alterations in RSFC are associated with greater severity of neurodevelopmental spectrum in middle childhood. This neurodevelopmental factor includes questions tapping into inattention, hyperactivity, clumsiness, repetitive behavior, and impulsivity. Neurodevelopmental symptoms are associated with deficits in cognitive abilities, including attentional processes.^12,47^ Previous work using dimensions examined here found that the neurodevelopmental spectrum showed the strongest associations with cognitive factors (i.e., fluid and crystalized intelligence) as compared to other hierarchical dimensions.^12^ In the current study, the identified connectivity alterations indicate that greater neurodevelopmental symptoms are associated with impairments in specific networks [DMN (4 findings), CON (3 findings), and Other (3 findings)]. Several of these networks implicated in attention to both internal and external stimuli (i.e., DMN and CON),^48^ as well as the ability to re-orient attention between internal thoughts and external attention.^26^ In support of this, research indicates that specifically diminished anticorrelation between DMN and task positive networks (e.g., CON) may contribute to attention difficulties,^27^ with impairments in connectivity between these networks associated with worse developmental trajectories for attention.^26^

Considering associations that replicated across two samples, neurodevelopmental symptoms were linked to alterations in DMN connectivity with 7 of 13 networks. Elevated neurodevelopmental problems were associated with lower within-network DMN connectivity—a relationship that was significantly stronger than the associations found for the p-factor—and other alterations involving the DMN, including smaller anti-correlation with the CON network, and lower connectivity with the ‘Other’ network. These observations are consistent with previous research linking DMN alterations in ADHD symptoms.^27,32,49–52^ Although speculative, these findings indicate an important role for the DMN and point to disruptions in DMN regulated functions such as mental simulations and attention to internal thoughts^23,53^ as potentially important contributors to neurodevelopmental symptoms.

The current study also found associations between higher neurodevelopmental symptoms and connectivity impairments in the CON network, a network associated with information integration and salience attribution.^31^ This is consistent with previous research that found evidence for altered CON connectivity in association with neurodevelopmental symptoms.^54–56^ Impaired anticorrelations between both CON and ‘Other’ and DAN and ‘Other’ networks may be reflective of impairments in neural functional integration that may contribute to neurodevelopmental symptoms. Also higher neurodevelopmental symptoms were associated with greater connectivity between the CON and VAN networks, which may be indicative of increased attention to external environmental stimuli, consistent with some previous research on inattentive symptoms.^35,47,57^ However, these hypotheses about the potential functional significance of RSFC alterations are speculative and in need of direct testing.

Aside from the single p-factor finding, all significant RSFC results were found specifically with the neurodevelopmental factor, but not with other dimensions. Present findings suggest that in middle childhood alterations in RSFC network alterations are fairly specific to the neurodevelopmental spectrum, compared to other forms of psychopathology that are often co-morbid with ADHD symptoms, including externalizing and internalizing symptoms. This perhaps points to the critical importance of the neurodevelopmental dimension in terms of neural functioning in middle childhood.

These results indicate that the neurodevelopmental symptoms trans-diagnostically related to ADHD, ASD, OCD, and learning disabilities are specifically associated with RSFC alterations.

Several limitations should be noted. The ABCD data used in this study are cross-sectional. Future studies in this sample could examine the association between these RSFC metrics and changes in neurodevelopmental problems over time to shed light on these associations. The psychopathology measure is limited to parent report and a single, albeit comprehensive, assessment system (CBCL). However, most of the results between RSFC alterations and the neurodevelopmental spectrum remained when accounting for parental psychopathology. Future research should examine whether present results are generalizable to child-reported psychopathology. Further, results were conducted using the Gordon parcellation and future research should examine the results using another parcellation definition (e.g., Schaefer parcellation^58^). The findings in this study were smaller than reported estimates in other smaller samples.^26,27^ This may be expected with a large, non-clinical, heterogeneous sample. Additionally, due to computational challenges in analysis and data sharing in datasets of this size, the current ABCD processed data release only includes parcel-based results. However, future ABCD releases on voxel-wise results will be utilized in subsequent research. Lastly, RSFC was collected using different scanning protocols across sites (e.g., different FIRMM software, different scanner manufacturer); therefore, all analyses controlled for these scanning differences.

In summary, the current study constitutes first steps in examining associations between RSFC and the hierarchical dimensional structure of psychopathology in middle childhood. We found evidence that RSFC connectivity alterations were largely specific neurodevelopmental spectrum, and the other dimensions of psychopathology (e.g., internalizing, externalizing) were not significantly associated with it, except for a weak association between p-factor and DMN. Associations with the neurodevelopmental dimensions—but not p-factor—remained when accounting for parental psychopathology. The finding that the neurodevelopmental dimension emerged as the only one robustly related to alterations in RSFC across two large sample highlights the importance of delineating this factor in neuroimaging studies of psychopathology dimensions to better understand neural underpinnings. This suggests that further research on connectivity alterations should target specific dimensions of psychopathology, although comprehensive assessment of psychopathology is essential to confirming specificity.

## Acknowledgments

Data used in the preparation of this article were obtained from the Adolescent Brain Cognitive Development (ABCD) Study (https://abcdstudy.org), held in the NIMH Data Archive (NDA). This is a multisite, longitudinal study designed to recruit more than 11,500 children age 9-10 and follow them over 10 years into early adulthood. The ABCD Study is supported by the National Institutes of Health and additional federal partners under award numbers U01DA041022, U01DA041025, U01DA041028, U01DA041048, U01DA041089, U01DA041093, U01DA041106, U01DA041117, U01DA041120, U01DA041134, U01DA041148, U01DA041156, U01DA041174, U24DA041123, and U24DA041147. A full list of supporters is available at https://abcdstudy.org/nih-collaborators. A listing of participating sites and a complete listing of the study investigators can be found at https://abcdstudy.org/principal-investigators.html. ABCD consortium investigators designed and implemented the study and/or provided data but did not necessarily participate in analysis or writing of this report. This manuscript reflects the views of the authors and may not reflect the opinions or views of the NIH or ABCD consortium investigators.

The ABCD data repository grows and changes over time. The ABCD data used in this report came from DOI 10.15154/1460410.

This work was supported by National Institute on Drug Abuse grant U01 DA041120 (DMB), National Institute of Health grants MH014677 (NRK).

